# Endogenous CRISPR arrays for scalable whole organism lineage tracing

**DOI:** 10.1101/501551

## Abstract

The last decade has seen a renewed appreciation of the central importance of cellular lineages to many questions in biology (especially organogenesis, stem cells and tumor biology). This has been driven in part by a renaissance in genetic clonal-labeling techniques. Recent approaches are based on accelerated mutation of DNA sequences, which can then be sequenced from individual cells to re-create a “phylogenetic” tree of cell lineage. However, current approaches depend on making transgenic alterations to the genome in question, which limit their application. Here, we introduce a new method which completely avoids the need for prior genetic engineering, by identifying endogenous CRISPR target arrays suitable for lineage analysis. In both mouse and zebrafish we identify the highest quality compact arrays as judged by equal base composition, 5’ G sequence, minimal likelihood of residing in the functional genome, minimal off targets and ease of amplification. We validate multiple high quality endogenous CRISPR arrays, demonstrating their utility for lineage tracing. Our technique thus can produce deep and broad lineages *in vivo*, while removing the dependence on genetic engineering, and also avoiding the need for single-cell analysis.

## Body

Development describes the process whereby a single totipotent zygotic cell transforms into a complex multicellular-organism. Defining the early patterns of cell division in developing organisms is of paramount importance to understand and ultimately control the mechanisms of cell fate decisions that impact on developmental, stem cell and cancer biology. The traditional method for defining the early patterns of cell division focused on fate mapping, which when performed at cellular resolution is called lineage tracing^1–4^.

Original methods for labeling cells depended on direct injection of a chosen cell early in development, with dyes or enzymes that would be retained in daughter cells over multiple rounds of division^5^. A major improvement was the introduction of genetic methods, which removed the need for physical manipulation of the embryo. These relied on a stochastic molecular event permanently activating expression of a marker, which would be clonally inherited by all daughter cells of the cell of origin (for example the LacZ transgene^6^, or GFP transgene^7^). These “single-label” methods however, could not analyze multiple clones in the same piece of tissue, and were subsequently superseded by the various “rainbow-label” techniques in which the engineered stochastic genetic events activated random combinations of different fluorescent proteins^8^, thus allowing the labeling of many different clones with multiple different colours (figure 1a).

**Figure 1:**
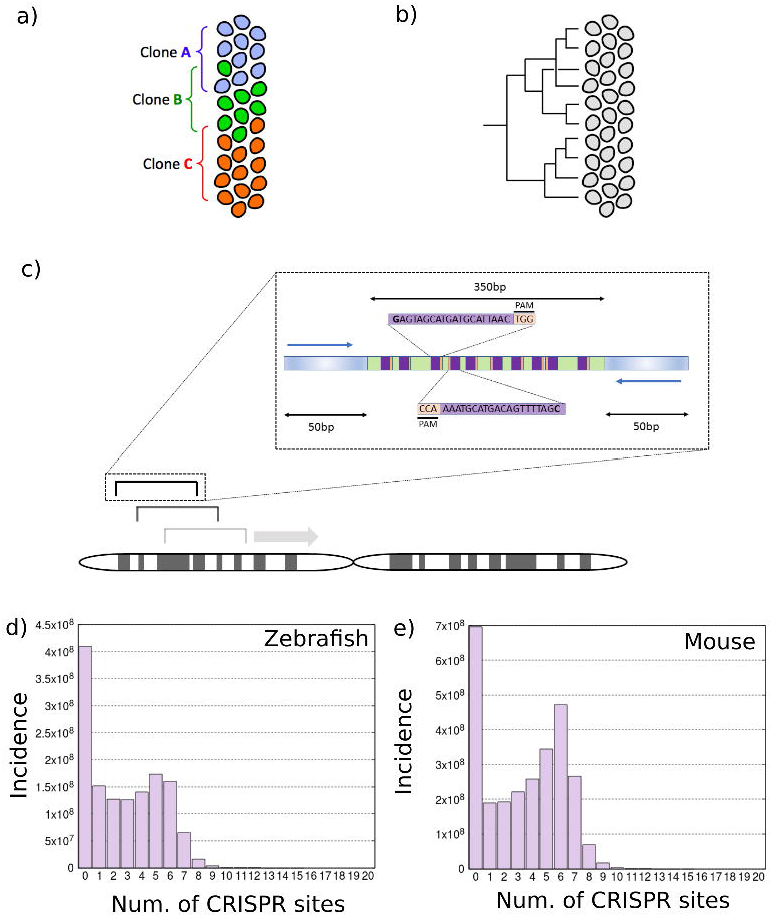
Identification of endogenous CRISPR arrays. a) Lineage dendrogram that can be generated with traditional genetic recombination approaches to lineage tracing. b) Lineage dendrogram that can be generated with whole-organism lineage tracing approaches. c) The Zebrafish and Mouse genomes were scanned using a moving window of 450bp. (inset) The flanking 50bp regions (shaded blue) are reserved for PCR amplification primer identification. We identify clusters of non-overlapping CRISPR targets within those windows. CRISPR target sites are illustrated by the purple box and the PAM sequences are shown in the red box. Primers are illustrated by the blue arrows. d and e) Histograms of the incidence (*y*-axis) of 350bp windows containing specific numbers of CRISPR target sites (*x*-axis) in the Zebrafish (d) and Mouse (e) genomes.

Clonal analysis distinguishes those cells within the clone from those outside, but involves only a single genetic event per clone, and thus does cannot contain rich information about the longer history of the cells. Any approach to re-create genuine cellular lineage trees from the end-point cells, requires the recording of multiple successive genetic changes within the same cell over time (figure 1b). The earliest attempts to use somatic mutation to generate cellular lineage trees focused on Microsatellite mutations that act as “molecular tumor clocks” that recorded past tumor histories^9^. This type approach using the genomic variability within an organism to elucidate the cell lineage tree has been described as “phylogenetic fate mapping”^10,11^.

This approach has been developed further to define cellular lineage relationships using genetically-engineered mice whose DNA-repair systems were compromised resulting in more mutations at the 120 MS loci analyzed^12^. Such mice have a high rate of mutations in MS loci, and develop a variety of spontaneous tumors^13^. This sped-up the accumulation of mutations, thus reducing the amount of sequencing required and allowing the first lineage trees to be achieved.

The recent advent of CRISPR technology^14^ has provided an alternative method for producing multiple independent mutations within the same cells. The targeted nature of CRISPR allows mutations to be targeted to a compact region of the genome called an array (instead of the 120 Microsatellites used in the Frumkin et al., 2008 for example) that can be readily deep sequenced. These approaches offer scalability to whole organism lineage tracing since each CRISPR target can potentially encode at least a single bit of information. Therefore the total amount of information encoded by an array of CRISPR sites is 2^n where n is the number of CRISPR target sites in the array. For lineage tracing the amount of information encoded by the array should be higher than the number of cells in the tissue that we want to lineage trace. To perform lineage tracing in the whole mouse embryo of 12 billion cells, 33 CRISPR target sites in an array (or multiple arrays) would be required to provide enough diversity.

Multiple approaches have employed CRISPR for lineage tracing *in vivo*. One such approach named GESTALT focuses on the generation of a synthetic compact array of CRISPR targets that was introduced into the genome of zebrafish^15^. Zygotic injection of the CRISPR machinery that targets that array therefore generates the diversity at that location which can be readily deep sequenced and used for lineage tracing. A second approach generates lineaging barcodes by targeting the same sequence in single or multiple repeats of a transgenic fluorescent protein genes^16–19^. However, both of these approaches suffer from the drawback of requiring the generation of a transgenic animal. Recently developed Tracerseq^20^ can be used on wildtype embryos but can only be used in conjunction with single cell sequencing platforms since it requires barcodes to be sequenced that have integrated into different regions of the genome.

Here we set-out to discover if a practical method of CRISPR-based lineage analysis could be achieved without having to genetically engineer the genome in advance. In particular, whether endogenous sites within the genome could act as suitable CRISPR arrays for this task. This approach would have the advantage of functioning on wildtype embryos simply by injecting the CRISPR machinery into the 1-cell zygote (or later stage). CRISPR target sites are constrained by the requirement for a proto-spacer adjacent motif (PAM) that has the form NGG for Cas9 (or CCN on the opposite DNA strand)^21–24^. Since a GG or CC dinucleotide is expected to arise on average every 8 base pairs, we reasoned that by chance arrays of compact CRISPR targets should appear naturally in most reasonably sized genomes. We explain the criteria employed to find suitable CRISPR arrays, illustrate our findings for zebrafish and mouse genomes, and validate in *in vitro* two of those arrays demonstrating that the target sites are efficiently edited as expected and that the method can indeed be used for lineage tracing.

## Results

To search for suitable endogenous CRISPR arrays we obtained appropriate genomic regions for Mouse and Zebrafish from the UCSC genome browser (see methods). We focused on regions of the genome that could be constructed with paired end reads on the Illumina Miseq since this platform offers an appropriate balance between paired-end read length (up to 2×150bp) with maximal throughput (number of reads). A small amount of overlap between the paired end reads allows for the region to be reconstructed and here we allow 50bp of overlap. Hence we searched the genome using a conservative 450bp moving window (figure 1c) which would allow efficient sequencing of clusters using a Miseq version 2 or 3 reagent kit. We reserve the first and last 50bp of the sequence for primer identification in order to amplify the region via PCR. This resulted in a window of 350bp for searching for the maximal number of CRISPR sites.

We searched for PAM sequences (NGG on the sense strand or NCC on the antisense strand) throughout the genomes of zebrafish and mouse. Overlapping target sites suffer from the drawback that editing events at one of the overlapping sites are likely to destroy the target site of another overlapping site, thus reducing the potential variability that can be generated in the CRISPR array. Therefore in order to maximize the variation amongst our CRISPR targets, we focused on non-overlapping CRISPR target sites (figure 1c inset) by searching for PAM sequences that had a space of at least 22 base pairs between them (Only 3bp between a GG followed by a CC as the target sequence would be read in opposite directions from the PAM). Histograms of the frequency of windows with different numbers of non-overlapping CRISPR targets sites with 5’G nucleotides are shown in figure 1d and e for the zebrafish and Mouse genomes respectively. As expected there are a huge number of windows with many CRISPR target sites across the genomes of the zebrafish and mice, as predicted from the frequency of CC and GG dinucleotides. There is a peak in window frequency for windows containing 5 non-overlapping target sites for the zebrafish genome and 6 non-overlapping target sites for the mouse genome (figure 1d and 1e respectively).

This high frequency of windows with many non-overlapping CRISPR target sites allowed us to use very stringent selection criteria so that we could focus on identifying the best possible CRISPR arrays. We therefore set up a series of selection criteria with the aim of applying them in order from the most stringent to the least stringent. We defined a CRISPR array as a contiguous region of the genome with >8 CRISPR sites per 350bp window. A G nucleotide at position 1 of the target sequence has been found to aid expression of the sgRNA from promoters such as U6 and T7 (Ran et al., 2013). Therefore we included the constraint that all CRISPR target sequences of a cluster must have a 5’ G nucleotide. These initial computationally rapid but stringent filters vastly reduced the number of potential arrays that we needed to assess with the following more computationally expensive filters.

We next applied four filters in the following order; 1) Balanced base frequency filter, 2) Minimal off-target filter, 3) non-functional site filter and 4) Filter for suitable PCR primers. We chose to focus on windows with a balanced base frequency as we deemed that the higher information content would remove the likelihood of secondary structure in the sgRNA (see discussion). Therefore we set a balanced base frequency filter such that the base with the highest frequency (count in window) could not be more than 50% of the base with the lowest frequency. We employed a filter that selects for endogenous CRISPR arrays with minimal off-targets because off targets potentially cause detrimental effects on the organism and also potentially quench the activity of Cas9 therefore leading to less efficient editing of our region of interest. Therefore, for each potential sgRNA target array we created a Bowtie2^25^ query file consisting of the set of target sequences. We then used Bowtie2 to search a prebuilt index of the corresponding genome. We set the Bowtie2 options so that only the 2 highest scoring alignments are reported for each sgRNA with sequences allowed to differ by up to 1 base pair. Arrays only passed the filter if no similar sequence could be found in the genome for any of the sgRNAs in the array. The non-functional site filter was used to restricted endogenous arrays to those that were less likely to have detrimental functional effects of the organism. We removed those that reside in a coding sequence or upstream regulatory region (up to 5kbp upstream). These are defined in the UCSC upstream5000 and mRNA files for mm10 and danrer10. We combined these putative functional sequences and built an index using bowtie2 for each species. We then used bowtie2 to search for the entire window of interest (450bp) in the functional sequences for that respective species. If the sequence was not found amongst the functional sequences then the array passed the filter. The final filter identified those CRISPR arrays that had flanking sequences with suitable primer binding sites so that the entire array could be amplified by PCR. We used Primer3^26^ to identify left and right primers in the 50bp flanking either side of the 350bp array window. If suitable left and right primers could be identified then the endogenous array passed the filter.

The results of the number of endogenous CRISPR arrays passing each filter for both species when we search for >8 CRISPR target sites are described with the nested Ven diagrams shown in figure 2a (see Supporting information tables 1 and 2 for corresponding values). The number of endogenous CRISPR arrays passing all filters can be seen to rapidly diminish when increasing the number of CRISPR target sites that are required per array. Interestingly all CRISPR arrays found using our criteria had suitable primer pairs for PCR amplification. The final distribution of endogenous CRISPR arrays over the genomes when searching for arrays with 9 CRISPR target sites is shown in figure 2b and c for zebrafish and Mouse respectively. As can be seen, potential CRISPR arrays exist on almost all chromosomes offering a flexible choice of targets. We provide a full list of the coordinates of endogenous CRISPR arrays (10 sites per array) for each species in bed format in the supporting information.

**Figure 2:**
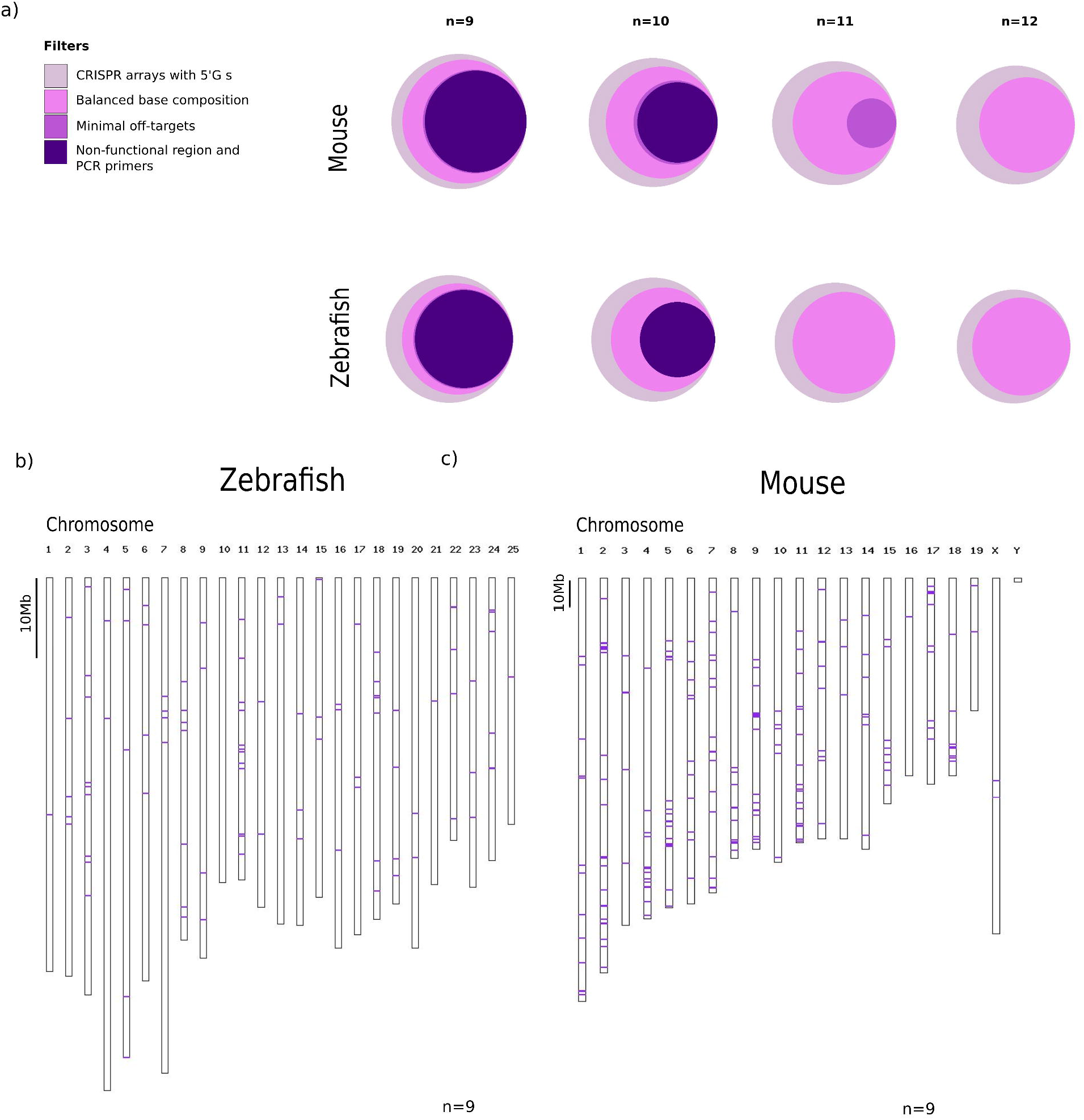
Summary of CRISPR cluster regions of the Zebrafish and Mouse genomes. a) Number of CRISPR arrays resulting from applying different filters in the Zebrafish and Mouse genomes. Areas represent the log of the number of arrays passing the filters in the nested Ven diagrams. b and c) Distribution of the filtered endogenous CRISPR arrays over the Zebrafish and Mouse genomes. Individual CRISPR arrays are represented by a red line. The result for 9 CRISPR targets per array is shown.

In order to validate our suggested target arrays, we assayed one array from each of the Mouse and Zebrafish genomes (M3 and Z4 - the oligonucleotide sequences to produce the corresponding sgRNAs are given in supporting information). We cloned the targeting sequences for the Mouse arrays into PX458^27^, which expresses both the sgRNA and Cas9 (Figure 3a). We transfected the resulting 10 mouse targeting vectors individually into mouse 3T3 cells and extracted genomic DNA 65 hours later. . For targeting in zebrafish we transcribed the 10 sgRNAs in vitro and individually microinjected them into the 1-cell stage zebrafish yolk sac with Cas9 protein (Figure 3b). Genomic DNA was extracted from embryos 30 hours later. We then amplified the M3 and Z4 arrays (shown with their spacer regions and PAM sequences in figure 3c and d respectively) and performed the surveyor nuclease assay.

The surveyor nuclease assay can detect the presence of mutations in a DNA fragment of interest if a wildtype reference exists. In the surveyor nuclease assay potentially mutated DNA is mixed with reference wildtype DNA. The two are melted and annealed resulting in hybrid double stranded DNA with bulges if a mutation was present. Surveyor nuclease is then added which cuts hybrid DNA at these bulges of miss-matched DNA. The resulting DNA is then run on a gel and two fragments are detected whose sizes correspond to the position of the miss-match relative to the ends of the DNA fragment if a mutation was present. CAS9 typically introduces indels in DNA 3-4bp upstream of the PAM site allowing us to predict the fragment sizes produced in the surveyor nuclease assay for each of our target sites. The expected fragment sizes depend on the location of the PAM site in the array. More peripheral PAM sites produce a small and a large fragment, whilst the most central PAM sites result in two similar sized fragments. The resulting spectrum of fragments resembles an X shape for both the M3 and Z4 arrays (figure 3e and f). The fragment sizes were then measured using a high sensitivity DNA chip and the results are shown in figure 3g and h. This result confirms that 19 out of 20 of our targets produce DNA fragments of the expected size (Only M3_10 has no discernable band). Therefore the endogenous CRISPR arrays that we have identified are genuine functional arrays that can be used for lineage tracing.

**Figure 3:**
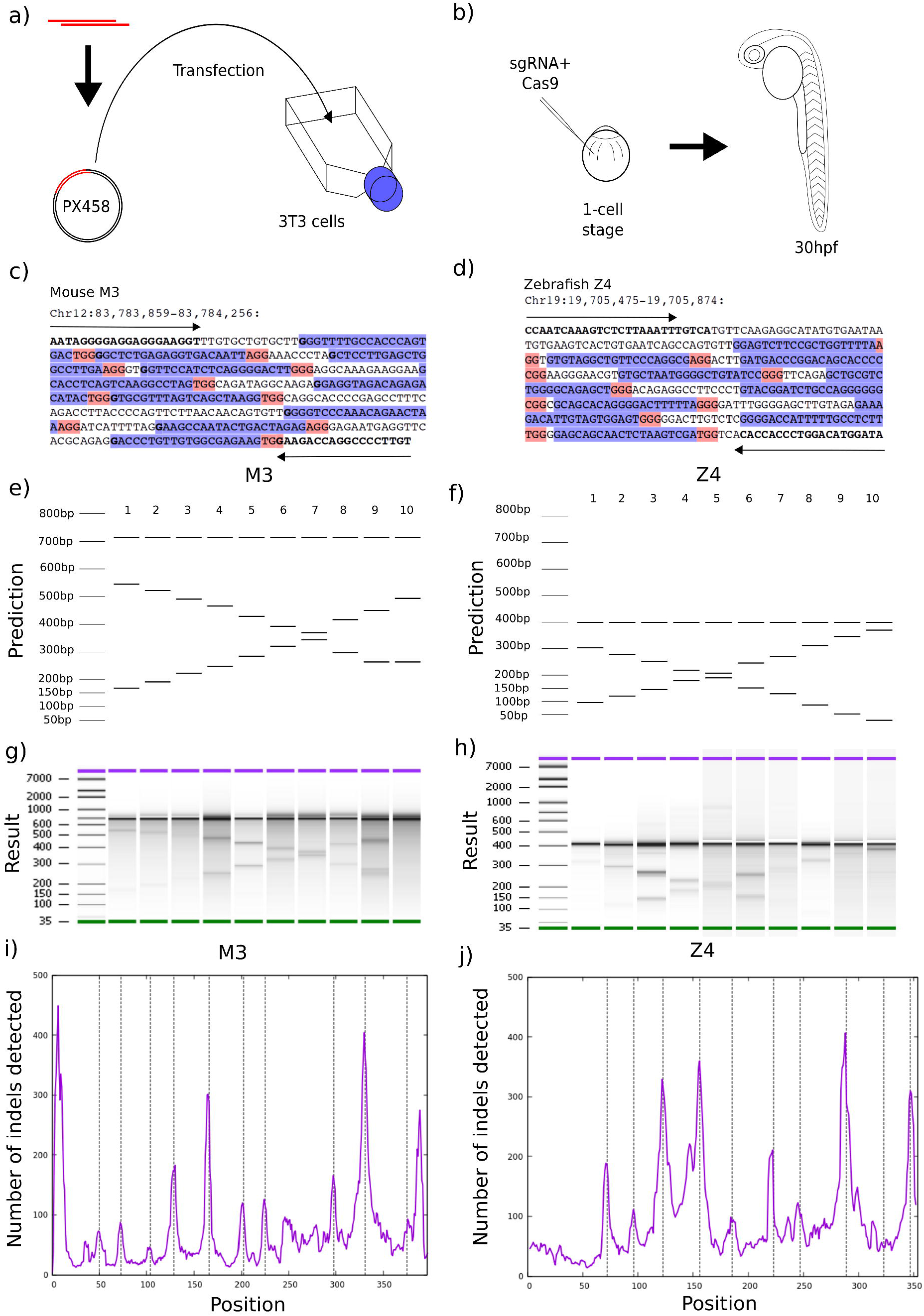
Validation of 2 target clusters from Mouse and Zebrafish. Protocols for validation of CRISPR targets *in vitro* in Mouse cell lines (a) or *in vivo* in Zebrafish embryos (b). Examples of endogenous CRISPR arrays from Mouse (c) and Zebrafish (d). The Primer3 PCR primer sequences and 5’Gs of target sites are shown in bold. The CRISPR targets sites are shown in Blue. The PAM sequences are highlighted in red. e and f) Fragment size predictions from surveyor nuclease assay are shown for the two tested endogenous CRISPR arrays. The band at the higher molecular weight is the uncut amplicon and the two bands at lower molecular weight are the cut fragments. g and h) Gels from a high sensitivity DNA chip after application of the surveyor nuclease assay. Bands are of the expected size for all targets with sufficient signal. i and j) Indel detection using Miseq deep sequencing of the pooled Mouse and Zebrafish amplicons. Purple lines show the number of indels (center point of indel and averaged over a window length of 5bp) detected at that specific position in the amplicon. Vertical dashed lines represent the expected positions of indels.

To further interrogate the editing ability of our targeting vectors we performed deep sequencing of the target arrays using the Miseq (Illumina) in order to quantify the frequency of insertion-deletion events (indels) over the arrays. We pooled the genomic DNA for all 10 samples used in the surveyor assay for both Mouse and Zebrafish. We then amplified the corresponding regions of the genome of these pools and performed deep sequencing using 2×250bp cycles on the Miseq (See methods). We identified the midpoint of each of the indels and calculated a histogram of editing incidence over the corresponding regions, The results are shown in figure 3i (M3), 3j (Z4) confirm that peaks of more frequent editing occur where we expect them (PAM position minus 4bp as represented by the dashed vertical lines). Furthermore we find an indel peak for M3_10 confirming that this targeting vector is also functional but that possibly the sensitivity of the surveyor nuclease assay is insufficient to detect it. Hence this result shows that indeed all 20 vectors are capable of editing their respective target site *in vitro*. Finally we picked two additional arrays for targeting each of the Mouse and Zebrafish genomes and assayed the CRISPR mutagenesis using illumina deep sequencing which further confirms that our lineaging arrays are functional (Figure S4).

In order to show that our tool could indeed be used for lineage tracing we microinjected 1-cell stage zebrafish embryos with CAS9 and an equimolar pool of the Z4 targeting sgRNAs and extracted genomic DNA 48 hours later (figure 4a). We then amplified the Z4 region and performed deep sequencing with the Miseq as described above. We focused on indels >=2 base pairs in size, as these were highly unlikely to result from sequencing errors. The spectrum of indels in each read was then used to construct a dendrogram using an inhouse program. In order to generate a representative metric of the utility of the approach we explored the dendrogram depth of all reads. The histogram of dendrogram depth for the Z4 array is shown in figure 4b showing that the full tree is up to 8 mutations in depth. The full dendrogram consists of 6,740 alleles and thus we show one sub-branch of the full dendrogram in figure 4c. A subset of that dendrogram is shown in figure 4d which demonstrates how reads are connected through the addition/removal of individual mutations. Taken together this data demonstrates how our approach can be used effectively to perform lineage tracing in cell lines.

**Figure 4:**
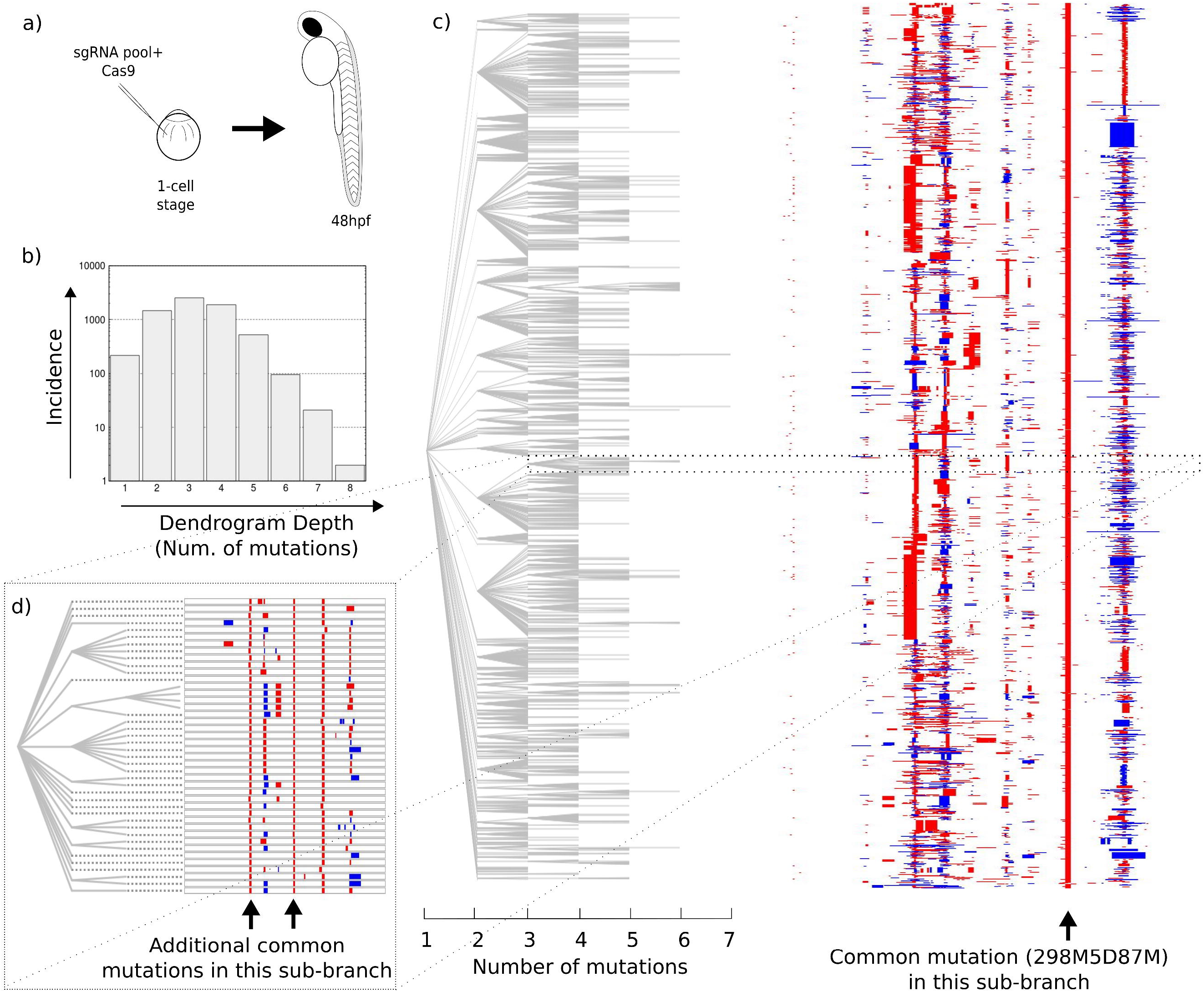
The approach can be used to perform lineage tracing. a) Schematic showing the approach to perform lineage tracing in the zebrafish. b) Histogram of the maximum depth of the reads in the dendrogram. c) Sub-branch of the dendrogram built using indel spectrum information of amplicons after deep sequencing (See Methods). Red blocks indicate a deletion and blue blocks indicate an insertion with the size of the block representing the size of the indel. d) One sub-branch of the dendrogram is highlighted in higher resolution.

## Discussion

By identifying endogenous CRISPR arrays with appropriate properties we demonstrate that it is not necessary to generate a transgenic animal to use CRISPR arrays for lineage tracing as in previous studies^15–19^. And importantly our technique also does not depend on single-cell sequencing. The approach described here can be easily extrapolated to any other species of interest. The proto-spacer adjacent motif (PAM) NGG is essential for CRISPR target sites. Given that a CC or GG dinucleotide is expected to occur on average every 8 base pairs, it is not surprising that we find a huge number of CRISPR target arrays throughout the zebrafish and mouse genomes. This large number of potential sites allows us to use strict selection criteria for optimal target site features resulting in the highest quality potential sites.

There are two important properties of any CRISPR based whole-organism lineaging system. Firstly the system must be capable of generating enough information (bits) to uniquely identify different cells. Here we show that our system is indeed capable of generating such diversity. The dendrogram shown in figure 4c is a single sub-branch of the full lineage dendrogram of this fish which consists of 6,740 alleles (Significantly higher than the 1,323 alleles identified in the V6 embryo from McKenna *et al*.,^15^). Secondly any whole-organism lineaging system must result in editing that is occurring throughout the duration of the developmental process so that multiple layers of a lineage tree can be defined. Given that our lineage tree has up to 8 levels (see figure 4b) our lineage system appears to be in tune with the developmental speed of the zebrafish embryo.

New technologies may expand the number of potential endogenous sites that can be used for whole organism lineage tracing. For example new nuclease enzymes that have different PAM sequence specificities such as cpf1, CasX and Y or Cas9 enzymes which have been engineered to alter their PAM specificities may increase the number of sites on offer^28–30^. The continued improvement in defining the most efficient sequences for targeting and minimizing off-targets also promises to refine our approach further^31–34^. Furthermore longer read sequencing technologies such as Pacbio and Oxford nanopore will result in deeper and broader lineaging dendrograms through the inclusion of more CRISPR sites and thus the generation of more bits of information. Other lineaging approaches have attempted to increase the number of available bits through self-targeting evolving barcodes^35,36^ but they have the drawback that it makes it more difficult to reverse engineer the history of the barcode and thus construct the lineage tree. Some groups have also recently used single cell sequencing alone at different developmental time points to reconstruct cellular lineage^37,38^.

The combination of a lineage tracing strategy such as that described here with single cell and spatial sequencing approaches will impact lineage tracing in multiple ways. Indeed, recently the use of single cell transcriptomics in combination with lineage tracing has been demonstrated in zebrafish to help uncover restrictions at the level of cell types, brain regions and gene expression cascades during differentiation^39^. Single cell sequencing will also allow multiple endogenous arrays located in different regions of the genome to be used for cell identification since it is then possible to discern which combination of array sequences came from the same cell (i.e. define the karyotypes). For example the Z3 and Z4 described here could be used together in combination, to generate deeper lineage trees. Though single cell sequencing approaches are currently much lower throughput that standard deep sequencing approaches they promise to improve in the future. Whole-organism lineage tracing in combination with single cell sequencing has also recently been used to aid in elucidating the mapping from progenitor cell to adult cell^18,20^,. Finally the combination of genetic lineaging approaches with techniques to define where cells are in physical space will become increasingly common. One such instance of performing CRISPR based lineaging tracing whilst defining where cells are in space with has recently been demonstrated *in vitro*^40^. We envision that all three of these techniques, genetic engineering-based lineaging tracing, spatial and single cell genomics/transcriptomics will be used combination to provide a rich plethora of information by which to address multiple questions in developmental and cancer biology.

Future versions of this system will involve inducible elements allowing lineaging to be addressed more effectively at later stages of development. Furthermore, prediction and measurement of sgRNA activity will be implemented when screening and selecting for endogenous arrays. One drawback of the current approach is that it is possible that both alleles from the same cell could be sequenced. Though this is likely to be rare, it could nevertheless lead to an erroneous lineage tree being generated. We envisage the use of Single Nucleotide Polymorphisms will circumvent this problem by giving information on which allele a read derives from. Therefore for each individual organism it is possible to generate 2 lineage dendrograms, one from each allele. Such internal repeats potentially offer greater reliability of lineaging since the dendrograms of each can be compared and coroborated.

To conclude, the advent of CRISPR technology has permitted the development of efficient whole organism lineage tracing tools. Our method described here is the first to use only endogenous CRISPR sequences from the wildtype genome, thus dramatically simplifying the procedure, and perhaps more importantly opening up the field of whole organism lineage tracing to non-model species that for which it is hard to generate transgenic animals.

## Methods

### Scanning the genome

We obtained the repeat masked genomes of Zebrafish (danrer10) and Mouse (mm10) from the University of California Santa Cruz (UCSC) genome browser. In order to focus on sequences whose genomic origin we could accurately identify and target we discounted the ChrUn, ChrM and ChrN_random data. Genomes were scanned using a moving window of 450bp. The first and last 50bp of each window was reserved for primer identification. The remaining central 350bp was analyzed for the presence of non-overlapping CRISPR target sites.

### Initial filters

At least 8 non-overlapping sites consisting of G(21xN)GG or CC(21xN)C had to be identified in order for the window to be processed further. All target sites had to have a 5’ G nucleotide for sense targets or a 3’C nucleotide for anti-sense targets. Base frequency was quantified in the window and a filter included that only allowed the processing of windows where the most frequent base was at most 50% more common than the least frequent base.

### Assaying off targets

Selected arrays were then further filtered by off target analysis. The set of potential CRISPR target sites (The 20bps corresponding to the targeting region) from each array was then queried with Bowtie2. Bowtie2 used a prebuilt index generated from the repeat masked Zebrafish and Mouse genomes (danrer10 and mm10 respectively). The important Bowtie2 flags used were N=1 and k=2. After calling bowtie we analyzed the number of alignment results. If that number was above the number of CRISPR sites queried then the array failed the filter since this means that there were at least 2 successful alignments for at least one of the CRISPR sites.

### Assaying for functional regions

Selected arrays were then assayed to see if they inhabited functional regions of the genome. We downloaded the mRNA and upstream5000 files of the zebrafish (Danrer10) and mouse (mm10) genomes from the UCSC genome browser. We combined these 2 files into one and constructed a bowtie2 index for each respective genome. We then called bowtie2 using the entire window of interest as a query sequence. The only Bowtie2 flag used was k=2. If the window did not align to anywhere in the function region then it passed the filter.

### Assaying whether the CRISPR array can be amplified

We passed the whole window of interest to a command line version of Primer3 with the tags:

SEQUENCE_TARGET=50,350,

PRIMER_PICK_LEFT_PRIMER=1, and

PRIMER_PICK_RIGHT_PRIMER=1

so that only primer sites on the flanking 50bp could be identified. Only if left and right primer sites in both the flanking regions could be identified did the window of interest pass the filter.

### Validation of CRISPR targeting in a Mouse cell line

Targeting vectors were generated by ligating annealed oligonucleotides corresponding to the sense and antisense of the target region into PX458 following the protocol of Ran et al., 2013. NIH3T3 cells were transfected with the appropriate plasmid and Lipofectamine 2000 according to the manufacturer’s instructions. 65 hours after transfection genomic DNA was extracted with the Qiagen Blood and Tissue kit. Genomic DNA was then further interrogated by surveyor Nuclease assay or Miseq deep sequencing as described below.

### Validation of CRISPR targeting and lineage tracing in Zebrafish

Targeting vectors were generated by annealing and extending the sense and antisense sgRNA targeting oligonucleotides given in supporting information using Phusion polymerase (98°C for 2 minutes, 50°C for 10 minutes and 72°C for 10 minutes). The NEB Hiscribe T7 transcription kit was then used to generate sgRNA (16 hours at 37°C) followed by the addition of DNase I for 15 minutes at 37°C. sgRNA was then purified using Zymogen clean and concentrator kit. sgRNA was microinjected at 100ng/µl into one-cell stage zebrafish embryo yolk sacs (AB and TL strains) with 8µM EnGen Cas9 (NEB) with 50 mM KCl, 3 mM MgCl_2_, 5 mM Tris HCl pH 8.0 and 0.05% phenol red. For validation of CRISPR targeting sgRNAs were microinjected individually and genomic DNA extracted 30 hours later using the Qiagen blood and tissue kit. For lineage tracing the Z4 sgRNAs were pooled and microinjected and genomic DNA extracted 48 hours later using the Qiagen blood and tissue kit.

### Surveyor Nuclease assay

Amplicons M3 and Z4 were amplified from genomic DNA using RedTaq polymerase (Sigma). For M3 we utilized an amplicon that was larger than our target array for this assay since this results in larger DNA fragments that are easier to detect. Due to imperfect transfection we expected amplicons to consist of a mix between wildtype (reference) sequences and mutated sequences. The hybrid mixes of amplicons were thus annealed by heating to 95°C and gradually ramping down to 25°C according to the surveyor nuclease protocol. Surveyor nuclease, enhancer and MgCl_2_ were then added to the hybridized PCR products in the volumetric ratio of 1:1:0.6:6. The surveyor nuclease reaction was then carried out at 42°C for 1 hour. The resulting reaction was stopped with 1/10 volume of STOP solution and the product cleaned using the Nucleospin PCR cleanup kit. DNA was eluted in 15µl of water and 1µl was run on a high sensitivity DNA chip (Agilent). Controls were run as suggested by the manufacturer.

### Miseq deep sequencing

We amplified the M3, M4, Z3 and Z4 amplicons with 2 rounds of PCR (20–35 and 15–20 cycles respectively) using the Q5 high fidelity polymerase (NEBNext 2x PCR master mix). The first round utilized the genomic DNA extracted from the 3T3 cells and zebrafish embryos using the standard M3, M4, Z3 and Z4 primers identified in our bioinformatic analysis. The second round was used to directly add the illumina adapters to these amplicons. Furthermore we added 5xN (random base pairs) directly 3’ to the Illumina universal primer in order to aid cluster resolution. Adapted amplicons were pooled and sequenced on the Illumina Miseq using the version2 sequencing kit at 2×250 cycles. Paired end reads were amalgamated using PEAR^41^ and the 5’ random 5xN bases were removed with cutadapt^42^. Amalgamated and trimmed reads were then mapped to the mouse genome (mm10) using bowtie2. We discarded reads which aligned perfectly to the genome (as these are unedited).

### Construction of dendrogram and quantification of dendrogram depth

We first defined a feature array for all reads by scanning through the CIGAR string of each mapped read and defining which particular mutation/combination of mutations occur within the 10 spacer regions. The spacer regions of the Z4 array used to generate the feature array were 81–101, 105–125, 132–152, 165–185, 194–214, 232–252, 256–276, 298–318, 332–352 and 356–376bp respectively. The exact position of the feature within the window was irrelevant which helps with offset differences between reads due to other mutations. We only considered indels >1bp to reduce the possibility that sequencing errors could add noise to the dendrogram.

We utilized an in-house program operating what we term a ‘catch’ method to construct the dendrogram and measure the depth where each read is located. The catch method starts with reads containing a single mutation and performs single-mutation steps bringing all reads into the catch which are identical to the current read but with one additional mutation (by scanning through the feature array). This process is repeated until no more reads can be brought into the catch. This catch method defines the parents and daughters of each read and the depth (number of mutations) at which each read is found (Used to generate the histogram in figure 4b). In our approach therefore internal nodes of the dendrogram are also reads.

Nodes of the dendrogram correspond to reads of the deep sequence data. Dendrograms are built for sub-trees rooted at each of the reads with a single mutation. An in-house algorithm calculates the coordinates of the nodes of the dendrogram. The *x*-value of nodes is assigned based on their depth in the catch method. The *y*-value for internal nodes in the midpoint of the daughter nodes. *y*-values are calculated by sweeping across the tree and iteratively setting the *y*-values of the leaf nodes (nodes with no daughters) with an incrementing value.

The algorithm decides the order of node processing using the topology of the tree as defined by the parent/daughter node data. For each node, the algorithm only uses the parent with the highest number of daughter nodes which prevents nodes linking to multiple parents in the dendrogram.

