## Supporting Information for "Endogenous CRISPR arrays for scalable whole organism lineage tracing"

### Values for Ven diagrams

### Mouse (Table 1)

| | 9 | 10 | 11 | 12 |
|---|---|---|---|---|
| **Suitable base composition** | 1928455 | 531075 | 178899 | 66578 |
| **All have 5'G** | 189272 | 19936 | 4362 | 1392 |
| **No off targets** | 4865 | 266 | 7 | 0 |
| **Not in functional site** | 3648 | 169 | 0 | 0 |
| **Suitable primers for amplification** | 3648 | 169 | 0 | 0 |

### Zebrafish (Table 2)

| | 9 | 10 | 11 | 12 |
|---|---|---|---|---|
| **Suitable base composition** | 393400 | 189817 | 86350 | 28578 |
| **All have 5'G** | 16786 | 5498 | 3618 | 2041 |
| **No off targets** | 2585 | 90 | 0 | 0 |
| **Not in functional site** | 2006 | 90 | 0 | 0 |
| **Suitable primers for amplification** | 2006 | 90 | 0 | 0 |

**Bed data**

Here we provide simple bed data for the endogenous Mouse and Zebrafish CRISPR arrays. This data consists of the chromosome and coordinates in the chromosome (i.e. bed fields chrom, chromStart and chromEnd). We provide the data for arrays containing 10 CRISPR sites. Note though the suitable windows that we have found often overlap. Therefore below we have combined overlapping sites into a single site. The coordinates of the raw overlapping sites can be found in the additional Mouse_sites.bed and Zebafish_sites.bed files.

**Mouse (10 CRISPR sites):**
**chr11 60183468 60183983**
**chr11 86021127 86021698**
**chr11 120184898 121035921**
**chr12 83783839 83784290**
**chr15 31342717 31343201**
**chr15 78212133 78212585**
**chr2 104152737 104153188**
**chr4 135466814 135467267**
**chr4 140037184 140037634**
**chr7 24759668 24760123**
**chr7 37571928 37572416**
**chr7 79554689 79555172**

**Zebrafish (10 CRISPR sites):**
**chr11 25561277 25561727**
**chr11 38579014 38579472**
**chr19 19705451 19705952**
**chr2 5837697 5838149**
**chr3 32303570 32304112**
**chr3 42422353 42422804**

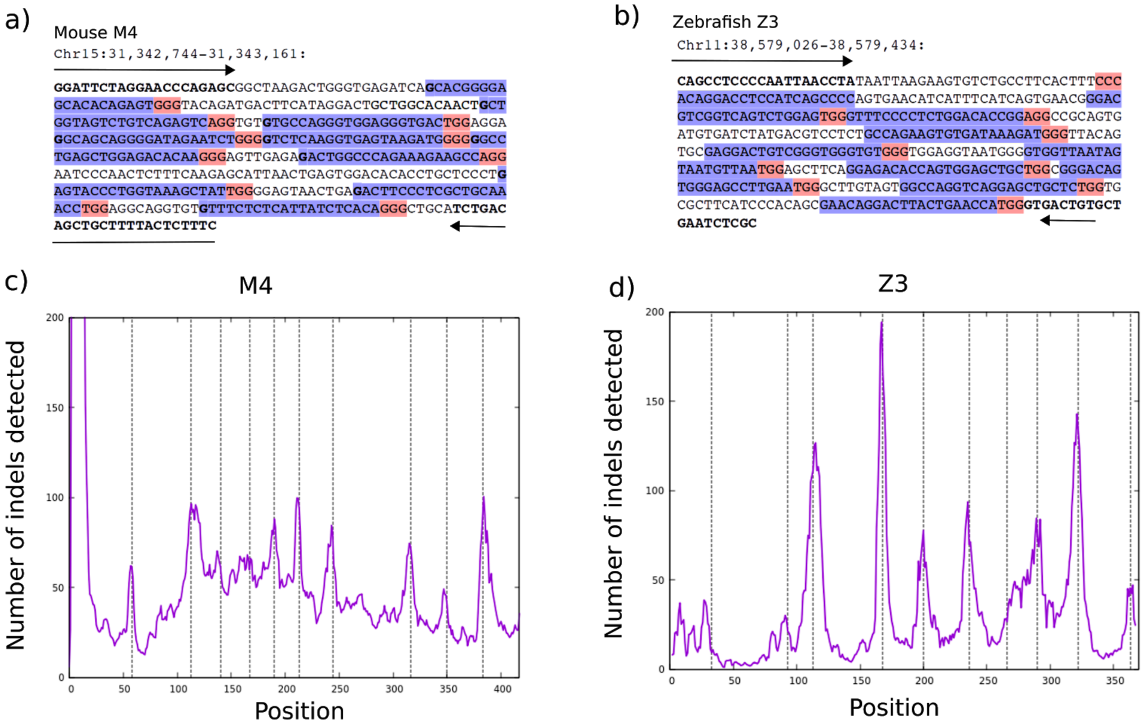


**Figure S1: Validation of additional endogenous CRISPR arrays from Mouse and Zebrafish.** a and b shows the arrays. Primer sites are show by arrows and bold text, spacer sites are highlighted in blue and PAM sites highlighted in red. s and d) Miseq deep sequencing demonstrated that the frequency of insertions and deletions have peaks at the expected editing locations (dashed lines).

**Oligonucleotides needed for generation of Zebrafish sgRNAs**

| Name | Sequence |
|---|---|
| **Universal_3prime** | AAAAGCACCGACTCGGTGCCACTTT TTCAAGTTGATAACGGACTAGCCTTA TTTTAACTTGCTATTTCTAGCTCTAAA AC |
| **Z3_1** | TTCTAATACGACTCACTATAG ACAGGACCTCCATCAGCCCC GTTTTAGAGCTAGAAATAGC |
| **Z3_2** | TTCTAATACGACTCACTATAG GGACGTCGGTCAGTCTGGAG GTTTTAGAGCTAGAAATAGC |
| **Z3_3** | TTCTAATACGACTCACTATAG GTTTCCCCTCTGGACACCGG GTTTTAGAGCTAGAAATAGC |
| **Z3_4** | TTCTAATACGACTCACTATAG GCCAGAAGTGTGATAAAGAT GTTTTAGAGCTAGAAATAGC |
| **Z3_5** | TTCTAATACGACTCACTATAG GAGGACTGTCGGGTGGGTGT GTTTTAGAGCTAGAAATAGC |
| **Z3_6** | TTCTAATACGACTCACTATAG GTGGTTAATAGTAATGTTAA GTTTTAGAGCTAGAAATAGC |
| **Z3_7** | TTCTAATACGACTCACTATAG GGAGACACCAGTGGAGCTGC GTTTTAGAGCTAGAAATAGC |
| **Z3_8** | TTCTAATACGACTCACTATAG GGGACAGTGGGAGCCTTGAA GTTTTAGAGCTAGAAATAGC |
| **Z3_9** | TTCTAATACGACTCACTATAG GGCCAGGTCAGGAGCTGCTC GTTTTAGAGCTAGAAATAGC |
| **Z3_10** | TTCTAATACGACTCACTATAG GAACAGGACTTACTGAACCA GTTTTAGAGCTAGAAATAGC |

| Name | Sequence |
| --- | --- |
| **Z4_1** | TTCTAATACGACTCACTATAG GGAGTCTTCCGCTGGTTTTA GTTTTAGAGCTAGAAATAGC |
| **Z4_2** | TTCTAATACGACTCACTATAG GTGTAGGCTGTTCCCAGGCG GTTTTAGAGCTAGAAATAGC |
| **Z4_3** | TTCTAATACGACTCACTATAG GATGACCCGGACAGCACCCC GTTTTAGAGCTAGAAATAGC |
| **Z4_4** | TTCTAATACGACTCACTATAG GTGCTAATGGGGCTGTATCC GTTTTAGAGCTAGAAATAGC |
| **Z4_5** | TTCTAATACGACTCACTATAG GCTGCGTCTGGGGCAGAGCT GTTTTAGAGCTAGAAATAGC |
| **Z4_6** | TTCTAATACGACTCACTATAG GTACGGATCTGCCAGGGGGG GTTTTAGAGCTAGAAATAGC |
| **Z4_7** | TTCTAATACGACTCACTATAG GCAGCACAGGGGACTTTTTA GTTTTAGAGCTAGAAATAGC |
| **Z4_8** | TTCTAATACGACTCACTATAG GAAAGACATTGTAGTGGAGT GTTTTAGAGCTAGAAATAGC |
| **Z4_9** | TTCTAATACGACTCACTATAG GGGGACCATTTTTGCCTCTT GTTTTAGAGCTAGAAATAGC |
| **Z4_10** | TTCTAATACGACTCACTATAG GAGCAGCAACTCTAAGTCGA GTTTTAGAGCTAGAAATAGC |

**Oligonucleotides needed for cloning Mouse sgRNA targeting sequences into PX458**

| Name | Sequence |
| --- | --- |
| M3_1_sense | |

| | |
|---|---|
| | CACCGGGGTTTTGCCACCCAGTGAC |
| M3_1_antisense | AAACGTCACTGGGTGGCAAAACCCC |
| M3_2_sense | CACCGGGCTCTGAGAGGTGACAATT |
| M3_2_antisense | AAACAATTGTCACCTCTCAGAGCCC |
| M3_3_sense | CACCGGCTCCTTGAGCTGGCCTTGA |
| M3_3_antisense | AAACTCAAGGCCAGCTCAAGGAGCC |
| M3_4_sense | CACCGGGTTCCATCTCAGGGGACTT |
| M3_4_antisense | AAACAAGTCCCCTGAGATGGAACCC |
| M3_5_sense | CACCGGCACCTCAGTCAAGGCCTAG |
| M3_5_antisense | AAACCTAGGCCTTGACTGAGGTGCC |
| M3_6_sense | CACCGGGAGGTAGACAGAGACATAC |
| M3_6_antisense | AAACGTATGTCTCTGTCTACCTCCC |
| M3_7_sense | CACCGGTGCGTTTAGTCAGCTAAGG |
| M3_7_antisense | AAACCCTTAGCTGACTAAACGCACC |
| M3_8_sense | CACCGGGGGTCCCAAACAGAACTAA |
| M3_8_antisense | AAACTTAGTTCTGTTTGGGACCCCC |
| M3_9_sense | CACCGGAAGCCAATACTGACTAGAG |
| M3_9_antisense | AAACCTCTAGTCAGTATTGGCTTCC |
| M3_10_sense | CACCGGACCCTGTTGTGGCGAGAAG |
| M3_10_antisense | AAACCTTCTCGCCACAACAGGGTCC |

| **Name** | **Sequence** |
|---|---|
| M4_1_sense | CACCGGCACGGGGAGCACACAGAGT |
| M4_1_antisense | AAACACTCTGTGTGCTCCCCGTGCC |
| M4_2_sense | CACCGGCTGGTAGTCTGTCAGAGTC |
| M4_2_antisense | AAACGACTCTGACAGACTACCAGCC |
| M4_3_sense | CACCGGTGCCAGGGTGGAGGGTGAC |

| | |
|---|---|
| M4_3_antisense | AAACGTCACCCTCCACCCTGGCACC |
| M4_4_sense | CACCGGGCAGCAGGGGATAGAATCT |
| M4_4_antisense | AAACAGATTCTATCCCCTGCTGCCC |
| M4_5_sense | CACCGGTCTCAAGGTGAGTAAGATG |
| M4_5_antisense | AAACCATCTTACTCACCTTGAGACC |
| M4_6_sense | CACCGGGCCTGAGCTGGAGACACAA |
| M4_6_antisense | AAACTTGTGTCTCCAGCTCAGGCCC |
| M4_7_sense | CACCGGACTGGCCCAGAAAGAAGCC |
| M4_7_antisense | AAACGGCTTCTTTCTGGGCCAGTCC |
| M4_8_sense | CACCGGAGTACCCTGGTAAAGCTAT |
| M4_8_antisense | AAACATAGCTTTACCAGGGTACTCC |
| M4_9_sense | CACCGGACTTCCCTCGCTGCAAACC |
| M4_9_antisense | AAACGGTTTGCAGCGAGGGAAGTCC |
| M4_10_sense | CACCGGTTTCTCTCATTATCTCACA |
| M4_10_antisense | AAACTGTGAGATAATGAGAGAAACC |

**Oligonucleotides required for region amplification in Mouse and Zebrafish**

| Name | Sequence |
|---|---|
| **Z3_Forward** | CAGCCTCCCCAATTAACCTA |
| **Z3_Reverse** | GCGAGATTCAGCACAGTCAC |
| **Z3_RegionExpand_Forward** | TGGGACCAAAACTTCTGTTTG |
| **Z3_RegionExpand_Reverse** | GCATATTAAGGCCCCGATTT |
| **Z4_Forward** | CCAATCAAAGTCTCTTAAATTTGTCA |
| **Z4_Reverse** | TATCCATGTCCAGGGTGGTG |
| **Z4_RegionExpand_Forward** | AACAAATGTCTTGCAGGACTGA |
| **Z4_RegionExpand_Reverse** | TGCTGTTTGGGATGTAACAA |
| **M3_Forward** | AATAGGGGAGGAGGGAAGGT |

| | |
|---|---|
| **M3_Reverse** | ACAAGGGGCCTGGTCTTC |
| **M3_RegionExpand_Forward** | CGTGGAACAGGAGCAGAAG |
| **M3_RegionExpand_Reverse** | TGAGGTGTGGCTAGAGACAGG |
| **M4_Forward** | GGATTCTAGGAACCCAGAGC |
| **M4_Reverse** | TGCAGCCCTGTGAGATAATG |
| **M4_RegionExpand_Forward** | CCTTTCAGTGCAGTAACTCATGC |
| **M4_RegionExpand_Reverse** | CCCACGAGGAAAAAGAAATG |

**<u>Oligonucleotides needed to append illumina adapters to amplicons</u>**

| Name | Sequence |
|---|---|
| Z3_Illumina_Uni_F | AATGATACGGCGACCACCGAGATCTAC ACTCTTTCCCTACACGACGCTC TTCCGATCTNNNNNCAGCCTCCCCAATT AACCTA |
| Z3_Illumina_IX_R<br>Where X is the illumina index number | CAAGCAGAAGACGGCATACGAGATXXX XXXGTGACTGGAGTTCAGACGTGTGCT CTTCCGATCTGCGAGATTCAGCACAGTC AC<br>Where X signifies the corresponding bases of the Illumina indexes |
| Z4_Illumina_Uni_F | AATGATACGGCGACCACCGAGATCTAC ACTCTTTCCCTACACGACGCTC TTCCGATCTNNNNNCCAATCAAAGTCTC TTAAATTTGTCA |
| Z4_Illumina_IX_R<br>Where X is the illumina index number | CAAGCAGAAGACGGCATACGAGATXXX XXXGTGACTGGAGTTCAGACGTGTGCT CTTCCGATCTTGCAGCCCTGTGAGATAA TG<br>Where X signifies the corresponding bases of the Illumina indexes |
| M3_Illumina_Uni_F | AATGATACGGCGACCACCGAGATCTAC ACTCTTTCCCTACACGACGCTC |

| | |
|---|---|
| | TTCCGATCTNNNNNAATAGGGGAGGAG GGAAGGT |
| M3_Illumina_IX_R<br>Where X is the illumina index number | CAAGCAGAAGACGGCATACGAGATXXX XXXGTGACTGGAGTTCAGACGTGTGCT CTTCCGATCTACAAGGGGCCTGGTCTTC<br>Where X signifies the corresponding bases of the Illumina indexes |
| M4_Illumina_Uni_F | AATGATACGGCGACCACCGAGATCTAC ACTCTTTCCCTACACGACGCTC TTCCGATCTNNNNNGGATTCTAGGAACC CAGAGC |
| M4_Illumina_IX_R<br>Where X is the illumina index number | CAAGCAGAAGACGGCATACGAGATXXX XXXGTGACTGGAGTTCAGACGTGTGCT CTTCCGATCT-TGCAGCCCTGTGAGATA ATG<br>Where X signifies the corresponding bases of the Illumina indexes |
